# A Population Genetics Theory for piRNA-regulated Transposable Elements

**DOI:** 10.1101/2022.07.05.498868

**Authors:** Siddharth S. Tomar, Aurélie Hua-Van, Arnaud Le Rouzic

**Affiliations:** Université Paris-Saclay, CNRS, IRD, UMR EGCE, Gif-sur-Yvette, France

**Keywords:** Transposition regulation, Model, Simulations, piRNA clusters, Trap model

## Abstract

Transposable elements (TEs) are self-reproducing selfish DNA sequences that can invade the genome of virtually all living species. Population genetics models have shown that TE copy numbers generally reach a limit, either because the transposition rate decreases with the number of copies (transposition regulation) or because TE copies are deleterious, and thus purged by natural selection. Yet, recent empirical discoveries suggest that TE regulation may mostly rely on piRNAs, which require a specific mutational event (the insertion of a TE copy in a piRNA cluster) to be activated — the so-called TE regulation ”trap model”. We derived new population genetics models accounting for this trap mechanism, and showed that the resulting equilibria differ substantially from previous expectations based on a transposition-selection equilibrium. We proposed three sub-models, depending on whether or not genomic TE copies and piRNA cluster TE copies are selectively neutral or deleterious, and we provide analytical expressions for maximum and equilibrium copy numbers, as well as cluster frequencies for all of them. In the full neutral model, the equilibrium is achieved when transposition is completely silenced, and this equilibrium does not depend on the transposition rate. When genomic TE copies are deleterious but not cluster TE copies, no long-term equilibrium is possible, and active TEs are eventually eliminated after an active incomplete invasion stage. When all TE copies are deleterious, a transposition-selection equilibrium exists, but the invasion dynamics is not monotonic, and the copy number peaks before decreasing. Mathematical predictions were in good agreement with numerical simulations, except when genetic drift and/or linkage disequilibrium dominates. Overall, the trap-model dynamics appeared to be substantially more stochastic and less repeatable than traditional regulation models.

## 1 Introduction

Transposable elements (TEs) are repeated sequences that accumulate in genomes and often constitute a substantial part of eukaryotic DNA. According to the canonical ”TE life cycle” model (Kidwell and Lisch 2001; Wallau et al. 2016), TE families are not maintained actively for a long time in genomes. TEs are the most active upon their arrival in a new genome (often involving a horizontal transfer, Gilbert and Feschotte 2018); their copy number increases up to a maximum, at which point transposition slows down. TE sequences are then progressively degraded and fragmented, accumulate substitutions, insertions, and deletions, up to being undetectable and not identifiable as such. The reasons why the total TE content, the TE families, and the number of copies per family vary substantially in the tree of life, even among close species, are far from being well-understood, which raises interesting challenges in comparative genomics.

TEs spread in genomes by replicative transposition, which ensures both the genomic increase in copy number and the invasion of populations across generations of sexual reproduction. They are often cited as a typical example of selfish DNA sequences, as they can spread without bringing any selective advantage to the host species, and could even be deleterious (Orgel and Crick 1980; Doolittle and Sapienza 1980). Even if an exponential amplification of a TE family could, in theory, lead to species extinction (Brookfield and Badge 1997; Arkhipova and Meselson 2005), empirical evidence rather suggests that TE invasion generally stops due to several (non-exclusive) physiological or evolutionary mechanisms, including selection, mutation, and regulation. Selection limits the TE spread whenever TE sequences are deleterious for the host species: individuals carrying fewer TE copies will be favored by natural selection, and will thus reproduce preferentially, which tends to decrease the number of TE copies at the next generation (Charlesworth and Charlesworth 1983; Lee 2022). The effect of mutations relies on the degradation of the protein-coding sequence of TEs, which decreases the amount of functional transposition machinery, and thus the transposition rate (Le Rouzic and Capy 2006). Even though TEs can be inactivated by regular genomic mutations, as any other DNA sequences, there exist documented mutational mechanisms that specifically target repeated sequences, such as repeat induced point mutations in fungi (Selker and Stevens 1985; Gladyshev 2017). Alternatively, substitutions or internal deletions in TEs could generate non-autonomous elements, able to use the transposition machinery without producing it, decreasing the transposition rate of autonomous copies (Hartl et al. 1992; Robillard et al. 2016).

Transposition regulation refers to any mechanism involved in the control of the transposition rate by the TE itself or by the host. There is a wide diversity of known transposition regulation mechanisms; some prevent epigenetically the transcription of the TE genes (Deniz et al. 2019), others target the TE transcripts (Adams et al. 1997), or act at the protein level (Lohe and Hartl 1996). Recently, the discovery of piRNA regulation systems have considerably improved and clarified our understanding of TE regulation (Brennecke et al. 2007; Malone and Hannon 2009; Zanni et al. 2013; Ozata et al. 2019). piRNA regulation seems to concern a wide range of metazoan species (Huang et al. 2021), and acts on TE expression through a series of complex mechanisms, which can be summarized by a simplified regulation scenario known as the ”trap model” (Bergman et al. 2006; Kofler 2019). In such a scenario, regulation is triggered by the insertion of a TE in specific ”trap” regions of the genome, the piRNA clusters. TE sequences inserted in piRNA clusters (thereafter TE cluster insertions) are transcribed into small regulating piRNAs, that are able to silence homologous mRNAs from the same TE family by recruiting proteins from the PIWI pathway.

Early models, starting from Charlesworth and Charlesworth (1983), assumed that the strength of regulation increases with the copy number. The transposition rate is then expected to drop progressively in the course of the TE invasion up to the point where transposition stops. In contrast, the PIWI regulation pathway displays unique features that may affect substantially the evolutionary dynamics of TE families: (i) it relies on a mutation-based mechanism, involving regulatory loci that may need several generations to appear (ii) the regulatory loci in the host genome segregate independently from the TE families and have their own evolutionary dynamics (the TE amplifies in a genetically-variable population, which is a mixture of permissive and repressive genetic backgrounds), and (iii) the regulation mechanism may not be strongly dependent on genomic copy number. The consequences of these unique features on the TE invasion dynamics are not totally clear yet. Individual-based stochastic simulations have shown that piRNA regulation is indeed capable of allowing a limited spread of TEs, compatible with the TE content of real genomes (Lu and Clark 2010; Kelleher et al. 2018). Kofler (2019) has shown, by simulation, that a major factor conditioning the TE success (in terms of copy number) was the size of the piRNA clusters, while the influence of the transposition rate was reduced. The dynamics of transposable elements when regulated by a trap model thus appear to differ substantially from the predictions of the traditional population genetics models.

With this paper, we extend the existing corpus of TE population genetics models by proposing a series of mathematical approximations for the dynamics of TE copy number when regulated by piRNA in a ”trap model” setting. We studied three scenarios, differing by whether or not TE copies induce a fitness cost when inserted in piRNA clusters and/or in other genomic locations: Scenario (i) neutral TEs, (ii) deleterious TEs and neutral TE cluster insertions, and (iii) deleterious TEs and deleterious TE cluster insertions. We showed that an equilibrium copy number could be achieved in scenarios (i) and (iii), while the TE family decays (after having invaded) in scenario (ii). We confirmed that the transposition rate does not condition the equilibrium number of copies in the neutral model (i), but it does when TEs are deleterious. Model (iii), in which both genomic and TE cluster insertions are deleterious, lead to a complex transposition-selection-regulation equilibrium (the ”transposition-selection cluster” balance in Kofler (2019)), in which regulatory alleles do not reach fixation and stabilize at an intermediate frequency. The robustness of these predictions was validated by numerical simulations.

## 2 Models and methods

### 2.1 Population genetic framework

Model setting and notation traces back to Charlesworth and Charlesworth (1983), who proposed to track the mean TE copy number *n* in a population through the difference equation:

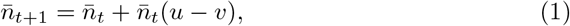

where *u* is the transposition rate (more exactly, the amplification rate per copy and per generation), and *v* the deletion rate. In this neutral model, if *u* and *v* are constant, the copy number dynamics is exponential. If the transposition rate *u_n_* is regulated by the copy number (*u*_0_ > *v*, d*u_n_*/d*n* < 0, and lim(*u_n_*) < *v*), then a stable equilibrium copy number 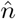 can be reached.

However, in most organisms, TEs are probably not neutral. If TEs are deleterious, fitness *w* decreases with the copy number (*w_n_* < *w*_0_). As a consequence, individuals carrying more copies reproduce less, which decreases the average copy number every generation. The effect of selection can be accounted for using traditional quantitative genetics, considering the number of copies *n* as a quantitative trait: 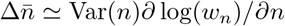, where Var(*n*) is the variance in copy number in the population, and ∂ log(*w_n_*)/*∂n* approximates the selection gradient on *n*. The approximation is better when the fitness function *w_n_* is smooth and the copy number *n* is not close to 0. Assuming random mating and no linkage disequilibrium, *n* is approximately Poisson-distributed in the population, and 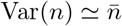.

Charlesworth and Charlesworth (1983) proposed to combine the effects of transposition and selection to approximate the variation in copy number among generations *t* and *t* + 1 as:

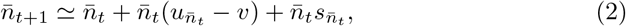

where 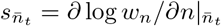.

When the transposition rate is high, the Poisson approximation does not hold and 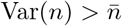: transposition overdisperses the copies in the population, as new TEs tend to appear in TE-rich genomes; random mating only halves this bias every generation but does not cancel it if transposition persists over generations. After transposition, the copy number rises to 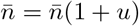, while its variance becomes 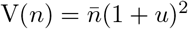; the drop in copy number due to selection thus becomes 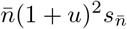 instead if 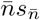. In order to match our simulation algorithm described below, in which selection takes place after transposition, we accounted for linkage disequilibrium and replaced the selection coefficient 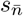 by 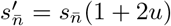 (neglecting *u*^2^ in (1 + *u*)^2^). This correction remains an approximation, as the effect of linkage disequilibrium of TE copies is more subtle and complex (Roze 2022). Overall, linkage disequilibrium due to transposition (slightly) amplifies the selection penalty for high transposition rates, and tends to decrease the genomic copy number.

### 2.2 Numerical methods

Data analysis was performed with R version 4.0 (R Core Team 2020). Mathematical model analysis involved packages deSolve (Soetaert et al. 2010) and phaseR (Grayling 2014). All figures and analyses can be reproduced from the scripts available at https://github.com/lerouzic/amodelTE.

Mathematical predictions were validated by individual-based simulations. Populations consisted in *N* = 1000 hermaphroditic diploid individuals, with an explicit genome of 30 chromosomes and a total of *K* = 10, 000 possible TE insertion sites (figure 1). *k* piRNA clusters of size *Kπ/k* were distributed on different chromosomes, the parameter *π* standing for the proportion of the *K* loci corresponding to piRNA clusters. Insertion sites were freely recombining, except within piRNA clusters. Generations were non-overlapping; reproduction consisted in generating and pairing randomly 2*N* haploid gametes from 2*N* parents sampled with replacement, with a probability proportional to their fitness. Transposition occurred with a rate *u_i_* computed for each individual as a function of its genotype at piRNA clusters (*u_i_* = *u* if no TE insertion in clusters, *u_i_* = 0 otherwise). The location of the transposed copy was drawn uniformly in the diploid genome. Transposition events in occupied loci were cancelled, which happened rarely as TE genome contents were always far from saturation (*K* ≫ *n*). Populations were initialized with 10 heterozygote insertions (in non-piRNA loci), randomly distributed in the population at frequency 0.05 each, resulting in *n*_0_ = 1 copy on average per diploid individual. For each parameter set, simulations were replicated 10 times, and the average number of diploid TE copies was reported. Average allele frequencies in clusters were calculated by dividing the number of diploid TE cluster insertions by 2k. The possibility to have two TE insertions in the same cluster was discarded, as such a situation requires two simultaneous transpositions in the same cluster: transposition is repressed in presence of TE cluster insertion, and there is no within-cluster recombination. We did not distinguish different regulatory alleles at the same cluster (i.e., TEs inserted in different sites within the cluster), as those are functionally equivalent. The simulation software was implemented in python (version 3.8.10 for Linux), with data structures from the numpy library (Harris et al. 2020). The code is available at https://github.com/siddharthst/Simulicron/tree/amodel.

**Figure 1:**
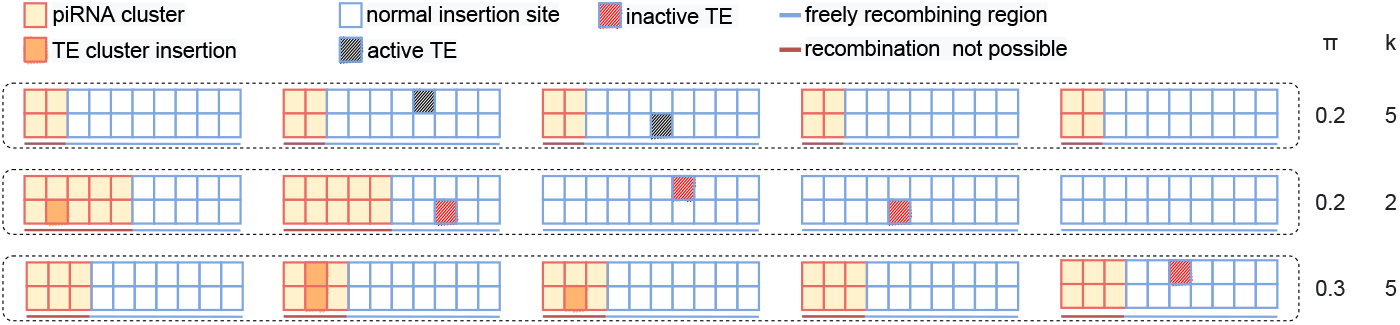
Schematic representation of the genomic model in the simulations. The *K* possible insertion sites are equally spread on chromosomes; the *k* piRNA clusters are distributed at one of the chromosome tips, representing a proportion *π* of the genome. Recombination is suppressed within clusters, but is possible in non-cluster genomic regions. In transposition-permissive genomes (no TE insertion in piRNA clusters), TEs located in normal insertion sites can transpose in any random location with a transposition rate *u*. TEs located in clusters cannot transpose, and prevent the transposition of all other elements in the genome. In order to ensure the generality of the results, simulations were set up to minimize genetic linkage: recombination between normal sites was free (recombination rate *r* = 0.5), and the genome had 30 chromosomes (always larger than the number of clusters). In the default conditions, 300 out of *K* = 10, 000 insertion sites are piRNA clusters (*π* = 0.03, close to the estimated proportion in the genome of *Drosophila melanogaster*).

## 3 Results

### 3.1 Neutral trap model

The model assumes *k* identical piRNA clusters in the genome, and the total probability to transpose in cluster regions is *π*. Each cluster locus can harbor two alleles: a regulatory allele (i.e., the cluster carries a TE insertion), which segregates at frequency *p*, and an ”empty” allele (frequency 1 – *p*). When all clusters are identical, and neglecting genetic drift (infinite population size), regulatory (TE cluster insertion) allele frequencies at all clusters are expected to be the same (*p*); at generation *t*, the average number of TE cluster insertions for a diploid individual is *m_t_* = 2*kp_t_*. TE deletions were neglected (*v* = 0). The model posits that the presence of a single regulatory allele at any cluster locus triggers complete regulation: the transposition rate per copy and per generation was *u* in ”permissive” genotypes (frequency (1 ‒ *p*)^2*k*^ in the population), and 0 in regulated genotypes (frequency 1 – (1 – *p*)^2*k*^). New regulatory alleles appear when a TE transposes in a piRNA cluster (with probability *π*), which is only possible in permissive genotypes. Assuming random mating and no linkage disequilibrium between clusters and the rest of the genome (Cov(*n_t_, p_t_*) = 0, i.e. no correlation between *n* and the genotype at the regulatory clusters), we approximated the discrete generation model with a continuous process, and the neutral model (equation 2) was rewritten as a set of two differential equations on 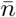 (relabelled *n* for simplicity) and *p*:

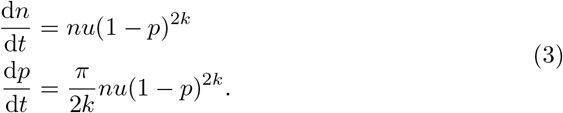

Here, *n* stands for non-cluster TE copies; for simplicity, equation 3 assumes that *π* ≪ 1, so that 1 – *π* ⋍ 1 (a more precise version of equation 3 for large values of *π* is provided in **Appendix A.1)**. Expressing the variation of the number of TE cluster insertions *m* = 2*kp* as d*m*/d*t* = *π*d*n*/d*t* would be more straightforward here, we kept the d*p*/d*t* formulation solely for consistency with more complex models described below.

The initial state of the system is *n*_0_ > 0 copies per individual in the population, and no piRNA cluster insertions (*p*_0_ = 0). The system of equation 3 admits three equilibria (characterized by the equilibrium values 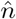 and 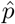): 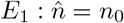 and 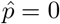 (no transposition, achieved when *u* = 0), 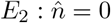 and 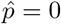 (no transposable element, *n*_0_ = 0), and 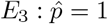 (fixation of all TE cluster insertions). Equilibria *E*_1_ and *E*_2_ do not need to be investigated further, as *u* = 0 or *n*_0_ = 0 do trivially result in the absence of TE invasion. Equilibrium *E*_3_ is analytically tractable, as d*n*/d*p* = 2*k/π*, and *n* = *n*_0_ + 2*pk/π* at any point of time:

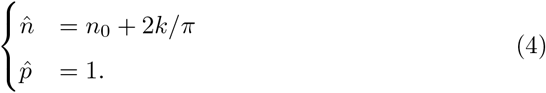

The fixation of regulatory alleles in all clusters is asymptotic (lim_*t*→∞_ *p* =1), and the equilibrium is asymptotically stable (d*n*/d*t* > 0 and d*p*/d*t* > 0). Figure 2 illustrates the effect of *u* and *k* on the dynamics *n_t_* and *p_t_*. The increase in regulatory allele frequency *p* is due to new transposition events in clusters (there would be multiple alleles at the sequence level, all being functionally equivalent). Assuming that *n*_0_ is small, the number of copies at equilibrium is proportional to the number of clusters *k*, and inversely proportional to the cluster size *π*. With several clusters (*k* > 1), the increase in copy number is slow, and the system may take a very long time to reach equilibrium. The copy number dynamics in simulations with different number of clusters remains very similar for the first hundreds of generations. The equilibrium copy number did not depend on the transposition rate *u*. This result relies on the absence of linkage disequilibrium between regulatory clusters and genomic copies; simulations showed that this assumption does not hold when the number of clusters increases, or when the transposition rate is very high (**Appendix B.1)**.

**Figure 2:**
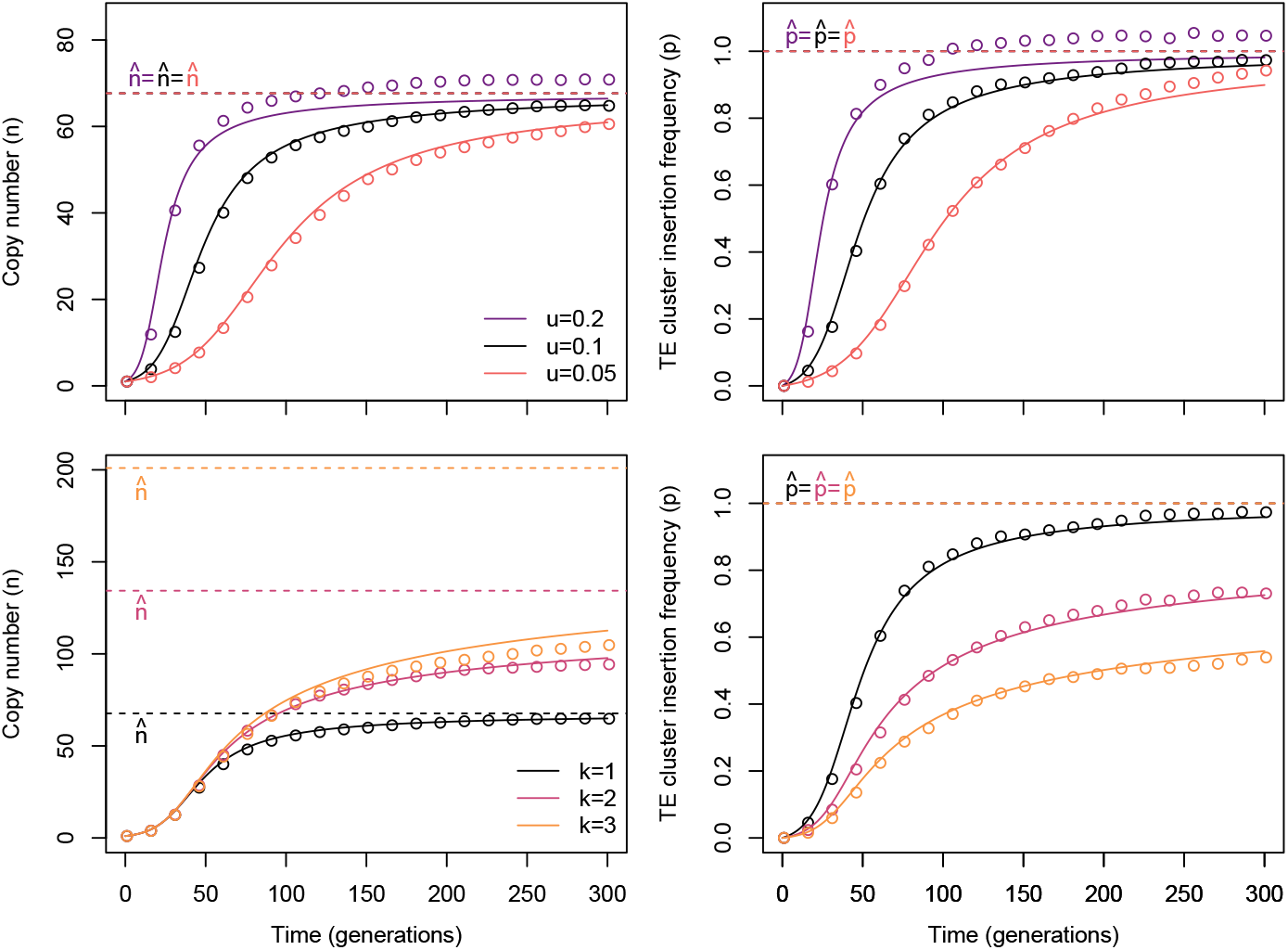
Dynamics in the neutral model. Unless indicated otherwise, default parameters were set to *n*_0_ = 1, *π* = 0.03, *k* = 1 cluster, and the transposition rate was *u* = 0.1. The top panel illustrates the influence of the transposition rate, the bottom panel of the number of clusters. Left: number of copies *n*, right: frequency of the TE cluster insertions in the population (*p*). Open symbols: simulations, plain lines: difference equations, hyphenated lines: predicted equilibria. The copy number equilibrium 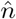 does not depend on the transposition rate, and the TE cluster insertion frequency at equilibrium 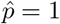 in all conditions. The time necessary to reach the equilibrium (both for 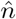 and 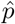) increases with *k*. In simulations, frequencies could be slightly > 1 due to rare events in which several TEs could insert simultaneously in the same cluster.

### 3.2 Selection

Natural selection, by favoring the reproduction of genotypes with fewer TE copies, generally acts in the same direction as regulation. A piRNA regulation model implementing selection could be derived by combining equations 2 and 3. In order to simplify the analysis, we derived the results assuming that the deleterious effects of TE copies were independent, i.e. *w_n_* = exp(–*ns*), where *n* is the copy number and *s* the coefficient of selection (deleterious effect per insertion), so that *∂* log *w_n_*/*∂n* = –*s*.

The following calculation relies on the additional assumption that piRNA clusters do not represent a large fraction of genomes (*π* ≪ 1, leading to *n* ≫ 2*kp*, i.e. that the number of TE copies in the clusters is never large enough to make a difference in the total TE count). We will describe two selection scenarios that happened to lead to qualitatively different outcome: TE insertions in piRNA clusters are neutral, and TE cluster insertions are as deleterious as the other insertions.

#### Deleterious TEs and neutral clusters

If TE cluster insertions are neutral, the model becomes:

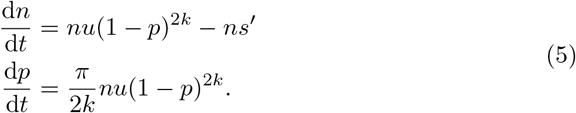

This equation only gives two equilibria, 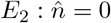 (loss of all copies), and 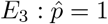 (when *s*′ = 0), which is the same as for the neutral model (equation 3): no selection and fixation of all TE cluster insertions. At the beginning of the dynamics, assuming *p*_0_ = 0, the TE invades if *u* > *s*′ (otherwise the system converges immediately to equilibrium *E*_2_ and the TE is lost). The copy number increases (d*n*/d*t* > 0) up to a maximum *n*\*, which is achieved when *p* = *p*\* (Figure 3). The maximum copy number can be obtained analytically (Appendix A.2):

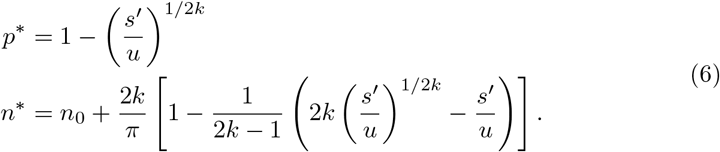

**Figure 3:**
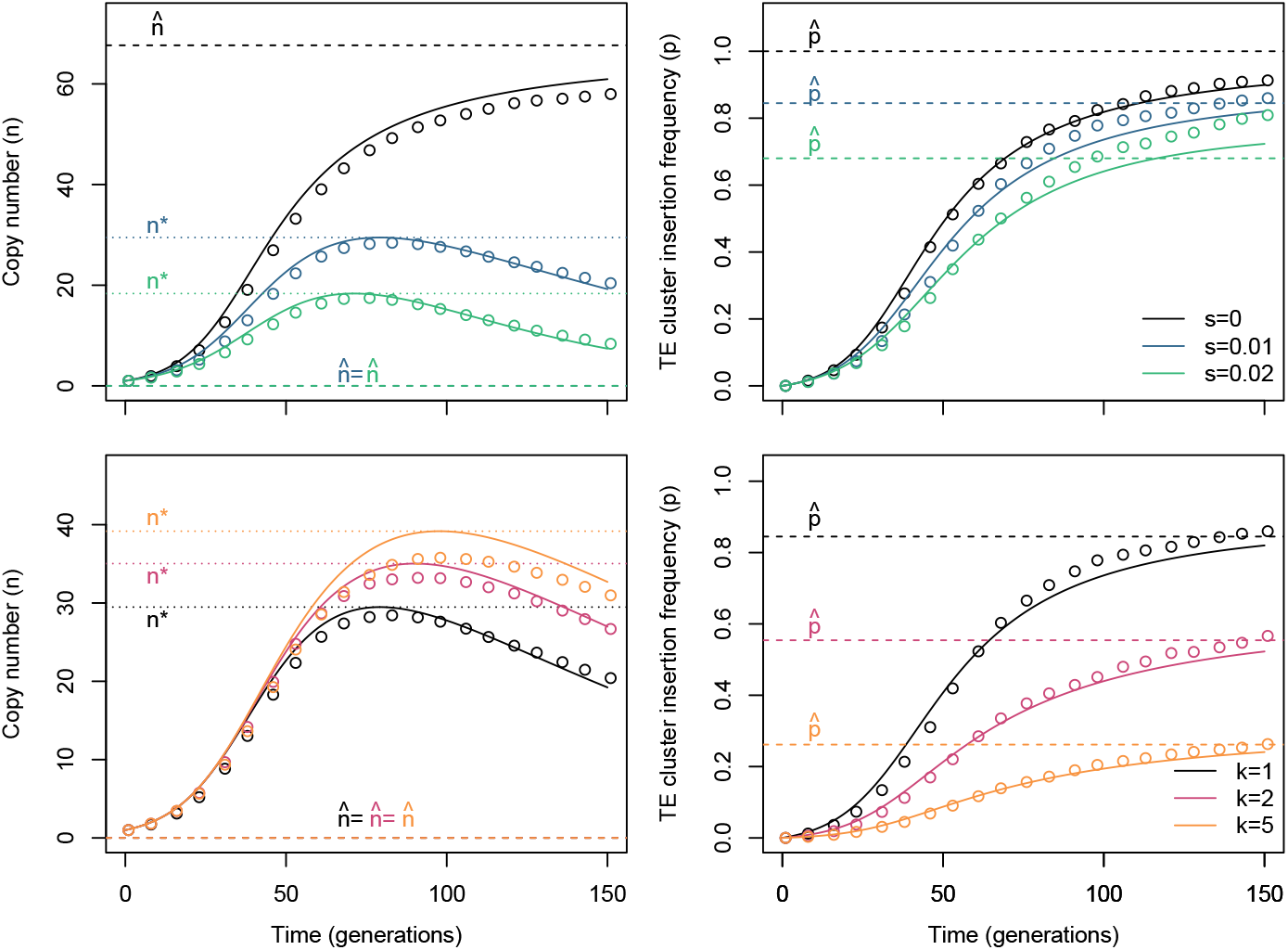
Dynamics of the deleterious TE - neutral cluster model. The top panel illustrates the influence of the selection coefficient *s* (with *u* = 0.1, *k* = 1), the bottom panel of the number of clusters *k* (with *s* = 0.01, *u* = 0.1). Left: number of copies *n*, right: frequency of the segregating TE cluster insertions in the population *p*. Open symbols: simulations, plain lines: difference equations, hyphenated lines: predicted equilibrium (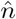 on the left, 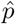 on the right), dotted lines: predicted copy number maximum *n*\*. Whenever *s* > 0, the copy number equilibrium 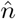 is 0.

The match between simulations (featuring a finite number of insertion sites, linkage disequilibrium, and non-overlapping generations) and this theoretical result was very good for a single cluster (*k* = 1), but degraded with larger number of clusters; *n*\* was overestimated by ~ 10% compared to simulations for *k* = 5 (Figure 3). Once the maximum number of copies is achieved, TE cluster insertions keep on accumulating, decreasing the transposition rate, which leads to a decrease in the copy number, up to the loss of the the element (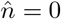 at equilibrium). At that stage, TE cluster insertions are not fixed, and the equilibrium TE cluster insertion frequency 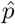 can be expressed as a function of copy number and TE cluster insertion frequency at the maximum (*p*\* and *n*\*) (Appendix A.3):

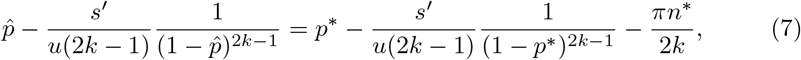

from which an exact solution for 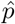 could not be calculated. The following approximation (from Appendix A.4):

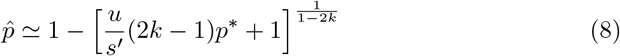

happens to be acceptable for a wide range of transposition rates and for small selection coefficients (*s* < 0.1) (Figure 4).

**Figure 4:**
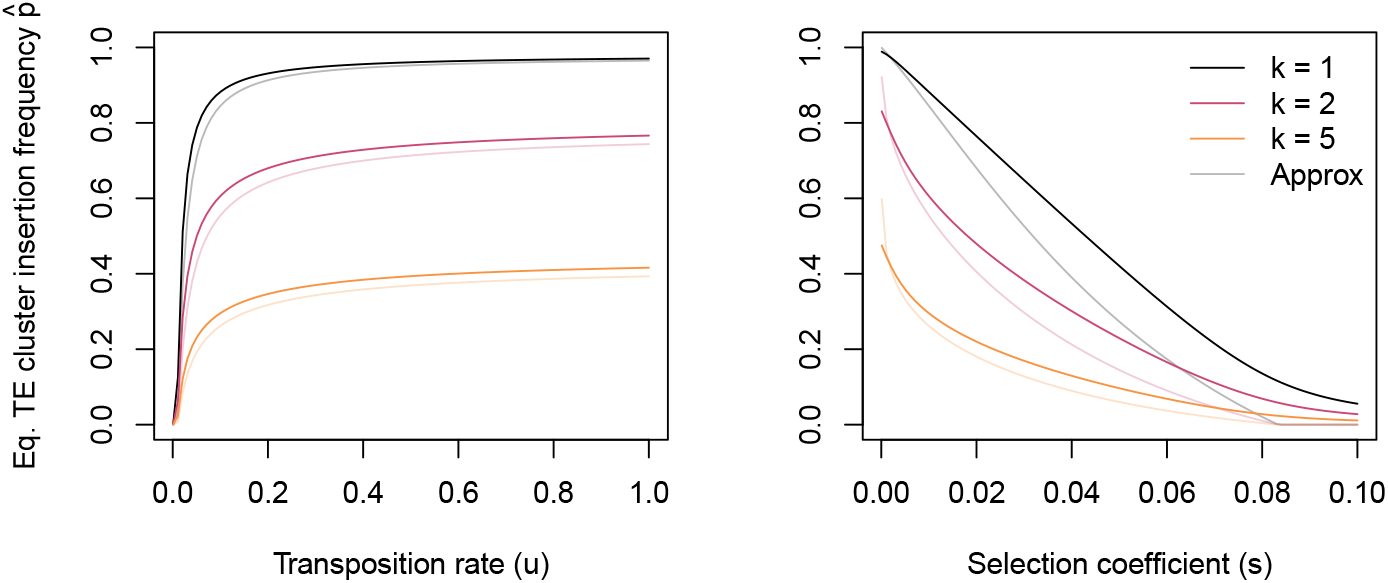
Influence of model parameters on the equilibrium TE cluster insertion frequency in the deleterious TE - neutral cluster model. The number of clusters *k* is indicated with different line colors. In this model, the TE is finally eliminated from the genome 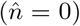, the equilibrium TE cluster insertion frequency 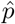 depends on the duration of the stay of the TE in the genome. Deleterious TEs are eliminated more rapidly, and have thus less opportunity to transpose into piRNA clusters, thus the lower 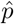. The approximation proposed in equation 8 is indicated in light colors.

The drop in 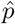 when the number of clusters *k* increases suggests that there might be a number of TE cluster insertions above which the transposition is effectively canceled. From equation 5, the copy number stabilizes when *u*(1 – *p*)^2*k*^ = *s*′, i.e. when *m* = 2*k*(1 – (*s*′/*u*)^1/2*k*^), where *m* = 2*kp* is the total number of TE cluster insertions per diploid individual. When TE copies are deleterious (*s* = 0.01), this expression tends to *m* = – log(*s*′/*u*) when *k* → ∞, which is about *m* = 2.1 copies per individual with our default parameters. In absence of selection, equation (4) predicts that transposition should occur up to the total fixation (i.e., *m* = 2*k*). However, the effective transposition rate will drop to very low levels much before fixation and its effects may rapidly become overwhelmed by genetic drift in finite-size populations. The variance of the change in copy number due to genetic drift between consecutive generations is about 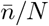; in a population of size *N* = 1000 with 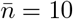 copies per individual, drift changes the average copy number by ±1%, while transposition will generate *u* exp(–*m*) new insertions in average (1% of 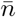 for *m* = 2 TE cluster insertions, and 0.2% of 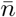 for *m* = 4 TE cluster insertions with our default parameter set *u* = 0.1). Based on simulations, Kofler (2019) estimated that transposition effectively stops above 4 TE cluster insertions per individual in average, which matches this prediction.

Equation 6 can be reorganized to address the problem of population extinction, as formulated in Kofler (2020). Numerical simulations have indeed shown that even if the final equilibrium state involves the loss of all TE copies, populations need to go through a stage where up to *n*\* deleterious copies are present in the genome. This makes it possible to approximate mathematically the critical cluster size *π*_c_, from which the population fitness drops below an arbitrary threshold *w_c_* and is at risk of extinction:

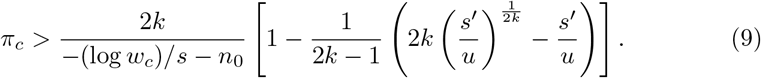

Setting *s* = 0.01, *u* = 0.1, and *n*_0_ = 1, as in the other examples, and taking *w_c_* = 0.1 gives *π_c_* > 0.0036 for *k* = 1 and *π_c_* > 0.005 for *k* = 5, these values being of the same order of magnitude than the interval 0.1% to 0.2% determined numerically by Kofler (2020).

#### Deleterious TEs and deleterious clusters

If the TE cluster insertions are as deleterious as other TEs, selection acts on TE cluster insertion frequency as predicted by population genetics (assuming no dominance):

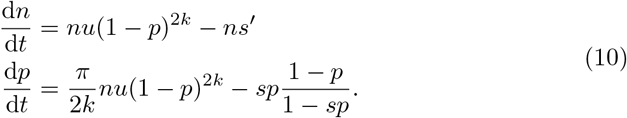

This allows for a new equilibrium *E*_4_:

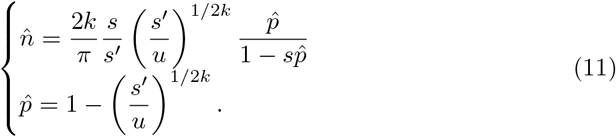

The equilibrium exists (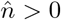 and 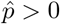) whenever *s* < *u*(1 + 2*u*), i.e. the transposition rate must be substantially larger than the selection coefficient. It corresponds to the ”Transposition-selection cluster” equilibrium described from numerical simulations in Kofler (2019). The dynamics for *n* and *p* are illustrated in Figure 5; the convergence to a non-zero equilibrium 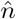 and an intermediate equilibrium for 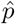 (no fixation) was confirmed by simulations. As for the neutral-cluster model, the match between simulations and mathematical predictions tends to degrade for large values of *k*. The influence of model parameters (*u, s*, and *π*) on equilibrium values are depicted in Figure 6; both the transposition rate *u* and the selection coefficient *s* show a non-monotonic relationship with the equilibrium copy number 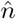 (maximum number of copies for an intermediate value of *u* and *s*).

**Figure 5:**
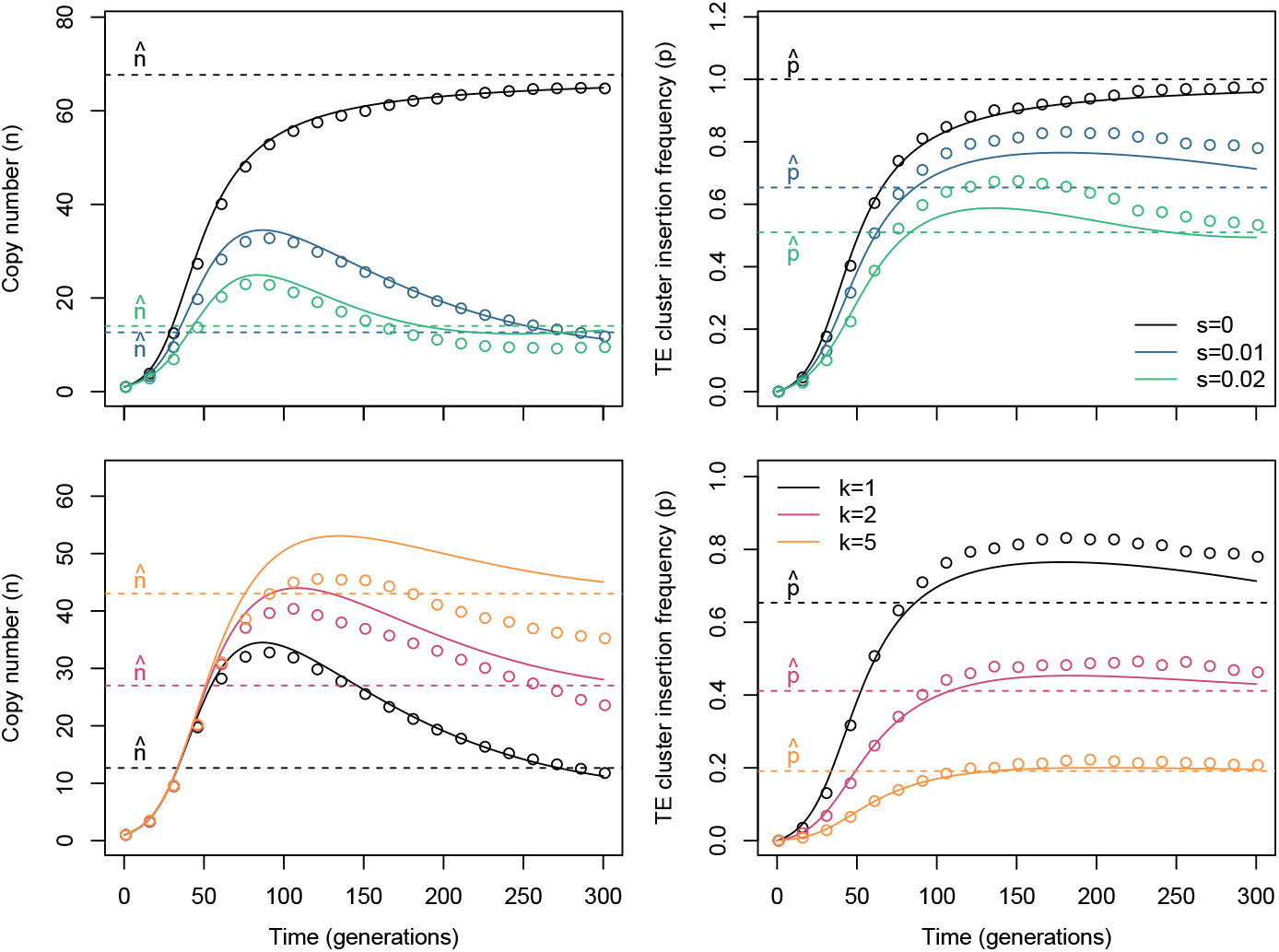
Dynamics of the deleterious TE - deleterious cluster model. Default parameters were *π* = 0.03, *k* = 1, *u* = 0.1, *s* = 0.01. Top panels: influence of the selection coefficient, bottom panels: influence of the number of clusters. Plain lines: predicted dynamics from equation 10, hyphenated lines: predicted equilibrium (eq. 11), open circles: simulations.

**Figure 6:**
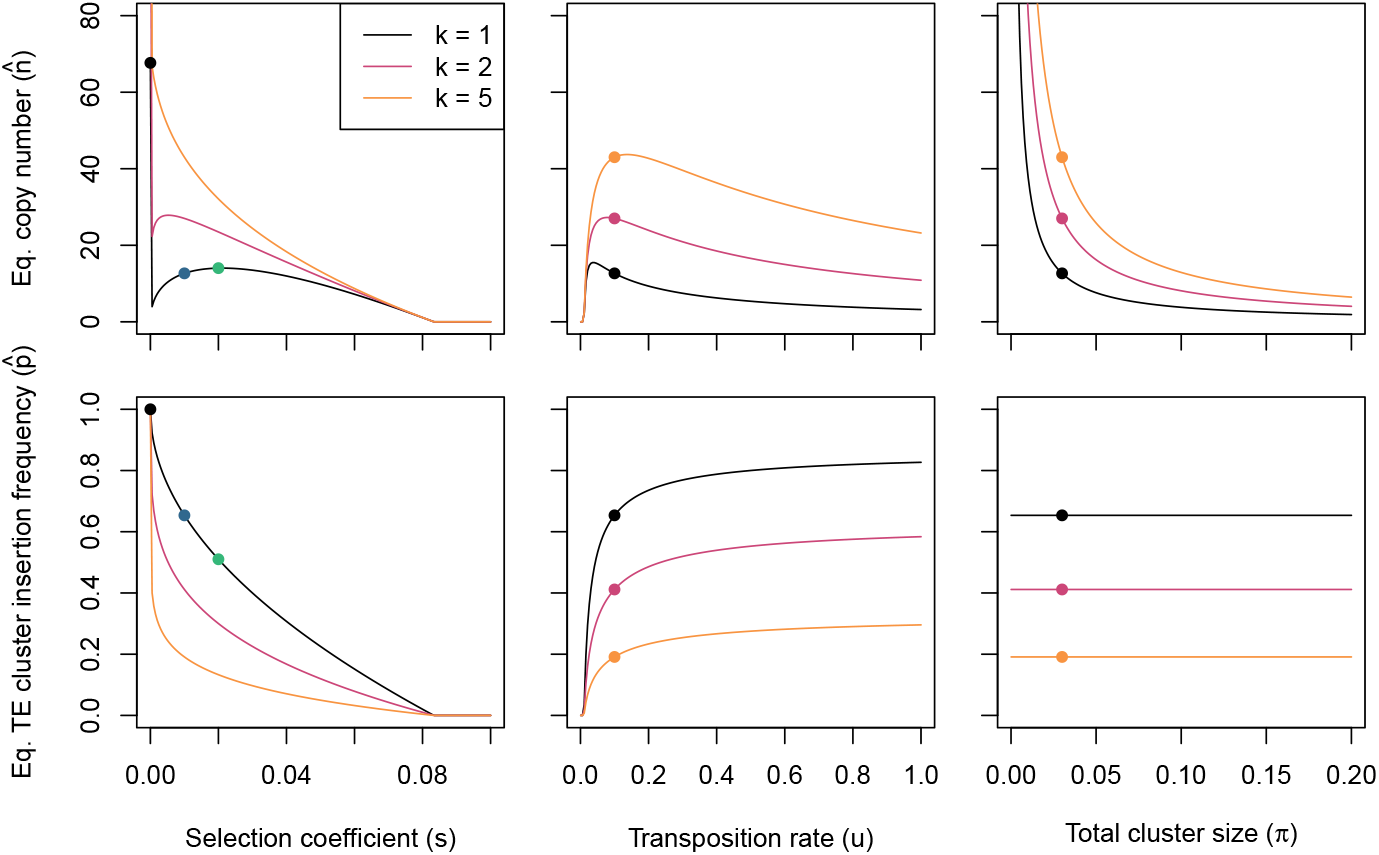
Influence of model parameters on the equilibrium TE copy number and the TE cluster insertion frequency in the deleterious TE - deleterious cluster model. Default parameter values were *u* = 0.1, *s* = 0.01, and *π* = 0.03. The number of clusters (*k* = 1, *k* = 2, and *k* = 5) is indicated by different colors. Colored dots indicate the equilibria illustrated in the panels of Figure 5 (same color code as in the figure).

A linear stability analysis (Appendix A.5) shows that for the whole range of *u, π*, and *k*, as well as for most of the reasonable values of *s*, the equilibrium is a stable focus, i.e. the system converges to the equilibrium while oscillating around it.

### 3.3 Genetic drift

The models described above assume infinite population size, which may not hold for low-census species and for laboratory (experimental evolution) populations. We assessed the influence of population size on the copy number with numerical simulations, comparing the neutral model, the deleterious TE - neutral cluster model, and the deleterious TE - deleterious cluster model with a ”classical” copy-number regulation model in which *u_n_* = *u*_0_/(1 + *bn*) (Charlesworth and Charlesworth 1983). Since the only equilibrium in the deleterious TE - neutral cluster model is the loss of all TE copies 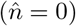, comparisons had to be performed at an intermediate time point of the dynamics (arbitrarily, after T = 100 generations). Models were parameterized such that the copy number *n* was approximately the same after 100 generations. Drift affects piRNA models substantially more than copy number regulation, the variance of all trap models being approximately one order of magnitude larger (Figure 7). Consistently with population genetics theory, the variance across simulation replicates decreased with 1/*N* for all models.

**Figure 7:**
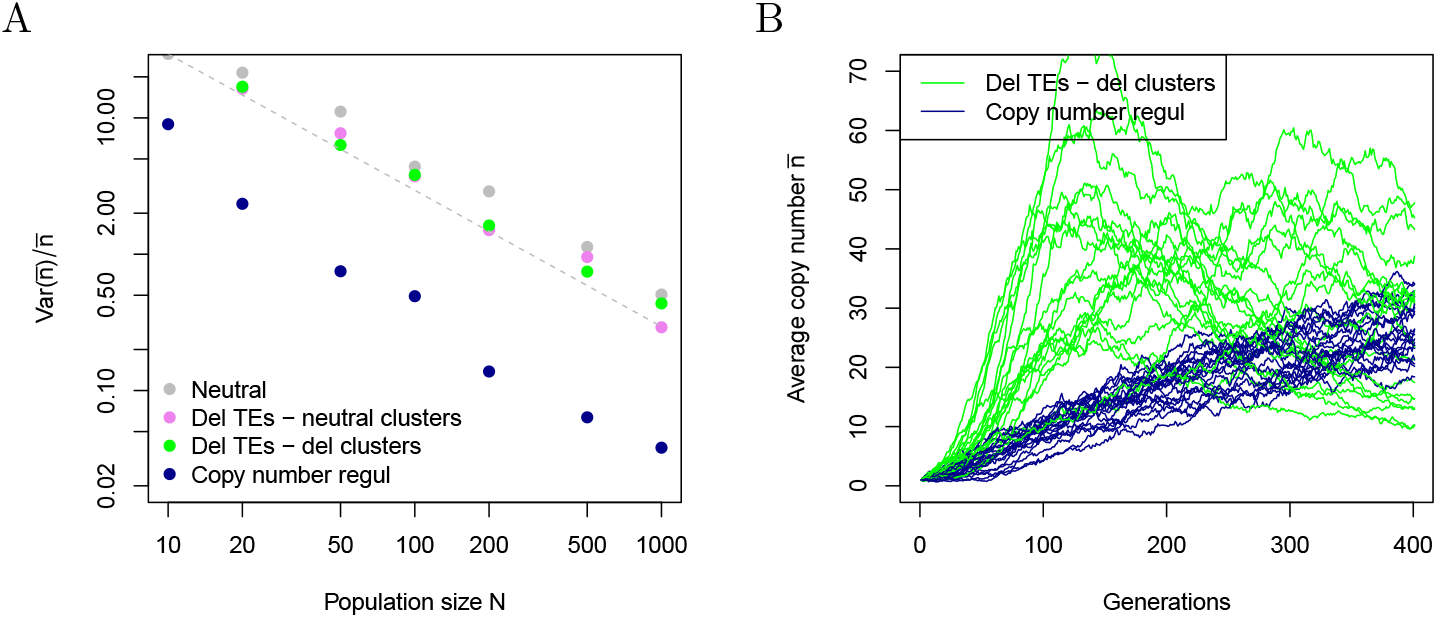
Effect of genetic drift on the TE copy number. A: Variance in the average copy number (relative to the average copy number) at generation 100 among replicated simulations for various population sizes. Four models are displayed: neutral trap model, deleterious TE - neutral clusters, deleterious TE - deleterious cluster, and copy number regulation. Models were parameterized so that they have very similar copy numbers (about 18) at generation 100; Neutral trap model: *u* = 0.045, *π* = 0.03, *k* = 2); Deleterious TE - neutral clusters: *u* = 0.13, *π* = 0.03, *s* = 0.01, *k* = 2; Deleterious TE - deleterious clusters: *u* = 0.07, *π* = 0.03, *s* = 0.01, *k* = 2; Copy number regulation: *u_n_* = 0.07/(1 + 0.3*n*), *s* = 0.01. The theoretically-expected decrease in variance (in 1/*N*) is illustrated for the neutral model (slope of –1 on the log-log plot). B: The effect of genetic drift is larger in the trap model than for copy-nuber regulation models. The figure displays the average copy number 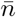 in 20 independent replicates, *N* = 100 for both models. Parameters were *u* = 0.1, *s* = 0.01, *π* = 0.03, *k* = 2 for the trap model, and *u_n_* = 0.1/(1 + 0.3*n*), and *s* = 0.01, for the copy-number regulation model. Regulation strengh was set so that the expected equilibrium copy number 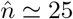 was the same for both models.

The standard population genetics theory predicts that selection in small populations is less effective at eliminating slightly deleterious alleles. Assuming that TE copies are deleterious, they should be eliminated faster in large populations compared to small ones. Although this mechanism has been proposed to explain the accumulation of junk DNA (including TE copies) in multicellular eukaryotes (Lynch and Conery 2003), little is known about how the equlibrium copy number of an active TE family is expected to be affected by drift even in the simplest scenarios (Charlesworth and Charlesworth 1983). Yet, informal models suggest that drift may have a limited effect, as copy number equilibria rely on the assumption that evolutionary forces that limit TE amplification (regulation and/or selection) increase in intensity when the copy number increases. Thus, when drift pushes the average copy number up or down, transposition is expected to be less or more effective respectively, which compensates the random deviation. Simulations show that, whatever the model, the copy number is indeed slightly higher in small populations (*N* < 100) when TEs are deleterious, but this effect never exceeds 20% of the total copy number (Supplementary figure B2). Overall, drift has a very limited impact on the mean copy number when *N* > 50.

## 4 Discussion

### 4.1 Population genetics of the trap model

The formalization of TE regulation by piRNA clusters (the ”trap model”) made it possible to derive a series of non-intuitive results, and evidence how trap regulation differs from traditional (copy-number based) regulation models. Among the most striking results: (i) in absence of selection (neutral trap model), the equilibrium copy number does not depend on the transposition rate, (ii) the proportion of genotypes able to regulate TEs increases with the size of clusters and decreases with the number of clusters, (iii) deleterious TEs can always invade when the transposition rate is larger than the selection coefficient, but the TE family can persist on the long-term only if TE cluster insertions are deleterious as well. When TE cluster insertions are neutral, they can increase in frequency up to fixation, which leads to the loss of all non-cluster TE copies. Equilibria are always stable. piRNA regulation being a mutational process, the TE copy number is more stochastic and substiantially more sensitive to genetic drift than other regulation models.

These results confirm and formalize previous work based on numerical simulations, in particular from Kofler (2019) who has already pointed out the small effect of transposition rate on the final state of the population and the inverse relationship between the number of clusters and the number of TE copies. The characterization of the equilibria demonstrate how the neutral trap model differs from the transposition-selection balance model proposed by Charlesworth and Charlesworth (1983); while the transposition-selection balance mostly depends on the transposition rate, the trap model equilibrium is determined by the mutational target (the size and the number of piRNA clusters).

While the equilibrium for the neutral trap model can be expressed with a very simple formula (equation 3), the derivation of copy number and TE cluster insertion frequencies is less straightforward when selection is accounted for (equations 10 and 11). When TEs are deleterious even when inserted in the clusters, the equilibrium copy number depends on the transposition rate *u* and the selection coefficient *s* in a non-monotonic way (less copies when *u* or *s* are either very low or very large). The fact that there exists an optimal transposition rate when TE insertions are deleterious have been proposed previously, in a different theoretical framework (Le Rouzic and Capy 2005). The elimination of high-transposition rate TEs from the genome is due to linkage disequilibrium; when transposition is very active, new TE copies are not randomly spread in the population but rather gathered into high-copy number (and thus, low fitness) genotypes. The optimal rate in the trap model (about 0.1 to 0.2 transpositions per copy and per generation in unregulated genetic backgrounds, figure 6) is compatible with empirical estimates (Robillard et al. 2016; Kofler et al. 2022).

### 4.2 Model approximations

The mathematical formulation of the trap model relies on a series of approximations. The general framework is strongly inspired from Charlesworth and Charlesworth 1983, and is based on the same general assumptions, such as a uniform transposition rates and selection coefficients among TE copies, diploid, random mating populations, and no linkage disequilibrium. This framework fits better some model species, including Drosophila or humans, than others (such as self-fertilizing plants or nematodes) for which the population genetics setup needs to be adapted. In general, individual-based (non-overlapping generations) simulations fit convincingly the predictions, but errors are cumulative in the trap model: small biases in the differential equations could add up over time and generate a visible discrepancy after several dozens generations.

The biology of the piRNA cluster regulation was also simplified. We considered that piRNA clusters were completely dominant and epistatic, i.e. a single genomic insertion drives the transposition rate to zero. Relaxing slightly this assumption is unlikely to modify qualitatively the model output, e.g. considering that regulatory insertions are recessive would change the frequency of permissive genotypes from (1 – *p*)^2*k*^ to (1 – *p*^2^)^*k*^, which would increase the TE cluster insertion frequency at equilibrium but not its stability. In a similar way, if the strength of regulation was increasing with the number of TE cluster insertions (or the number of genomic TEs), equilibrium would still be achieved for a higher number of copies. In contrast, imperfect regulation (a residual transposition rate even when all TE cluster insertions are fixed, such as in Lu and Clark 2010) would break the equilibrium in the neutral case, and copy number would raise indefinitely. This only affects the neutral model though, as imperfect regulation would have a much more limited effect when TEs are deleterious.

In order to compute the equilibria, we assumed no epistasis on fitness, i.e. constant *∂* log *w/∂n* = –*s*. Deriving the model with a different fitness function is possible, although solving the differential equations could be more challenging. Instead of our fitness function *w_n_* = *e^-ns^,* Charlesworth and Charlesworth (1983) proposed *w_n_* = 1 – *sn^c^* (*c* being a coefficient quantifying the amount of epistasis on fitness), while Dolgin and Charlesworth (2006) later used *w_n_* = *e*^-*sn-cn*2^ (different parameterizations for directional epistasis are discussed in e.g. Le Rouzic 2014). Considering negative epistasis on fitness (i.e. the cost of additional deleterious mutations increases) in TE population genetic models has two major consequences: (i) the strength of selection increasing with the copy number, it ensures and stabilizes the equilibrium even in absence of regulation (Charlesworth and Charlesworth 1983), and (ii) the model is more realistic, as epistasis on fitness for deleterious mutations has been measured repeatedly on many organisms (Maisnier-Patin et al. 2005; Kouyos et al. 2007; Khan et al. 2011). Interestingly, there is little evidence of negative epistasis for fitness among TE insertions (Lee 2022), suggesting that epistasis is probably not a major explanation for the stabilization of the copy number. In the trap model, regulation itself is strong enough to achieve an equilibrium in absence of selection, so epistasis on fitness is expected to modify the equilibrium copy number and the range of parameters for which a reasonable copy number can be maintained (Kofler 2019), but not the presence of a theoretical equilibrium.

Recent data might suggest that piRNA regulation may not be the only mechanism involved in early regulation of TE activity. For instance, Kofler et al. (2022) observed, in a lab experimental evolution context, that the number of TE cluster insertions was lower than expected in the trap model. Combining a copy-number regulation component and the trap model framework is theoretically possible and does not invalidate our approach, at the cost of introducing a new regulation parameter in the equations. In a more general way, the diversity of transposition regulation mechanisms in animals (Lu and Clark 2010; Saint-Leandre et al. 2020), plants (Roessler et al. 2018), and micro-organisms (Sousa et al. 2013), makes it impossible to derive models that are both accurate and universal.

### 4.3 piRNA clusters, selection, and recombination

The most effective configuration for TE regulation is a single, large piRNA cluster. Dividing the cluster in smaller parts increases equilibrium TE copy numbers, and reducing the total cluster size as well. Kofler (2019) has already noticed that recombination among cluster loci reduces the efficiency of regulation, and that regulation was more efficient with one large, non-recombining cluster than with many small clusters spread on several chromosomes (the single-cluster model was called the ”flamenco” model in Kofler 2019, inspired from the *flamenco* regulatory locus in Drosophila, Goriaux et al. 2014). In our setting, the proportion of transposition-permissive genotypes in the population is (1 – *m*/2*k*)^2*k*^ when *m* = 2*kp* TE cluster insertions are present in the genome and evenly distributed among clusters. This proportion increases as a function of *k*, meaning that the most efficient regulation is achieved when *k* = 1. Furthermore, the number of copies at equilibrium is expected to be proportional to the number of clusters *k*. Selection for TE regulation should thus minimize recombination within and across clusters; the fact that, in most organisms, piRNA clusters seem to be located at several loci needs to be explained by other factors (such as functional constraints) than the regulation efficiency. For instance, spreading piRNA clusters at several genomic locations may prevent TEs to evolve cluster avoidance strategies. The need to regulate independently different TE families might also play a role in the scattering of piRNA clusters; the interactions between several TE families invading simultaneously may generate new constraints on the regulation system, which probably deserves further investigation.

Our theoretical analysis displays contrasted results depending on the selection pressure on TE insertions located in piRNA clusters. The nature and the size of TE-induced deleterious effects is a long-standing issue (Nuzhdin 2000; Lee 2022) that remains poorly understood. TEs can affect the host fitness through various mechanisms, including the interruption of functional genomic sequences, chromosomal rearangements due to ectopic recombinations between TEs inserted at different loci, or a direct poisoning effect of the transposition process. The properties of piRNA clusters seem to limit most of these potential effects: the density of functional regions in probably low; clusters tend to be located in low-recombination regions, and the transcription into mRNA is likely to be repressed. It is thus tempting to speculate that the effect of TE cluster insertions on the host fitness should be lower than TE copies inserted in random genomic positions, and that the deleterious TE - neutral cluster model is more realistic than the deleterious TE - deleterious cluster model. Solid empirical evidence is necessary to confirm this speculation, which cannot be addressed solely based on theoretical considerations.

An interesting hypothesis was raised by Kelleher et al. (2018) about the possibility that TE cluster insertion frequency could be influenced by positive selection. Assuming deleterious TEs, genotypes able to control TE spread are indeed expected to display a selective advantage over those in which transposition is unregulated, suggesting that TE cluster insertions should sweep in the population as advantageous alleles. Our model, neglecting linkage disequilibrium between TEs and TE cluster insertions, would then underestimate the increase in frequency of regulatory alleles (and thus overestimate the copy number). Although the reasoning is theoretically valid, the actual strength of positive selection on piRNA clusters is probably limited in general. Assuming that TE insertions have a local deleterious effect (because they disrupt genes or gene regulation), the selective advantage of a regulatory locus is weak and indirect (of the same order of magnitude as *n* × *u* × *s*, the deleterious effect of the few insertions arising in a single generation). In contrast, if active transposition is deleterious (such as in the hybrid dysgenesis scenario explored by Kelleher et al. 2018), the selective advantage of TE cluster insertions is of the order of magnitude of *n* × *s*, and selection may have an effect on TE cluster insertion frequencies. Although it is experimentally difficult to determine how selection acts on TEs, both scenarios are expected to leave different genomic footprints, as the positive selection hypothesis posits that regulatory alleles should be shared among many individuals of the population, while the neutral hypothesis expects that various individuals are regulated by independent TE cluster insertions. Empirical evidence is scarce, but seems to favor the neutral hypothesis (Zhang et al. 2020).

### 4.4 Concluding remarks

Comparative genomics applied to transposable elements is hard. Notwithstanding the countless potential artefacts associated with sequencing, assembly, and annotation biases, understanding the evolutionary history of genomes is limited by the small number of evolutionary replicates, and the number of TE families and TE copy numbers accumulated in a single species is frequently dominated by stochastic and contingent factors. Being able to compare observed patterns with model predictions is thus of utter importance.

By extending the existing theory of transposable elements population genetics, we were able to demonstrate that the trap regulation model was affecting deeply the dynamics of TE invasion. In particular, when TE cluster insertions are neutral, the possibility to maintain a stable copy number equilibrium disappears, and all active TE copies are expected to be lost eventually. When regulatory insertions are slightly deleterious, a hypothetical transposition-selection equilibrium can be achieved, in which genomic TEs maintain as selfish DNA sequences, while regulatory insertions maintain as a result of a selection-mutation balance. This situation prevents the fixation of TE cluster insertions, and thus maintains a low frequency of transposition-permissive genotypes (without piRNA against active TE families), which could be measured empirically in populations to estimate the likelihood of the alternative regulation scenarios.

## Acknowledgements

The authors thank three anonymous reviewers for their positive and constructive suggestions.

## Appendix A Mathematical details

### A.1 Equation 3 for large *π*

Equation 3 assumes for simplicity that 1 – *π* ≃ 1, which may not hold in all species. If only a proportion 1 – *π* of new insertions fall in non-cluster regions, the model can be re-written as:

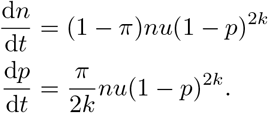

The equilibrium then becomes:

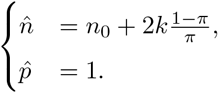

### A.2 Equation 6

When the copy number *n* achieves its maximum *n*\*, d*n*/d*t* = 0. This happens when the TE cluster insertion frequency *p*\* is:

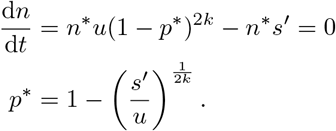

The number of copies cumulated while *p* is rising from *p*_0_ to *p*\* can be calculated by integrating both sides:

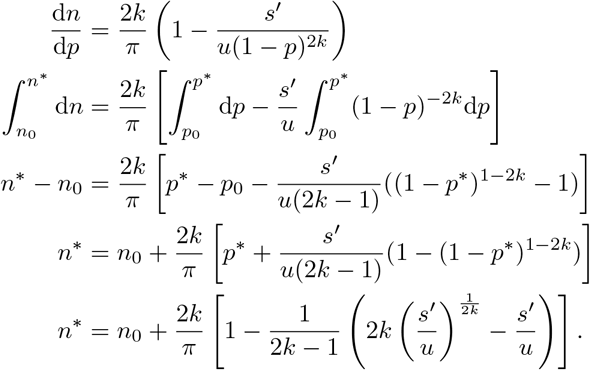

### A.3 Equation 7

The strategy was very similar than for obtaining *n*\*, with d*p*/d*n* integrated both sides from the maximum to the equilibrium:

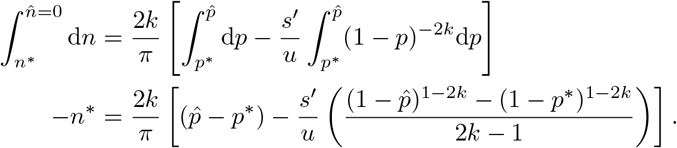

### A.4 Equation 8

Rewriting the previous equation with 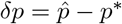 and 1 – *p*\* = *q*\* gives:

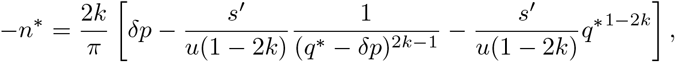

which turns out to be dominated by the second term (1/(*q*\* – *δp*)^2*k*-1^ ≫ *δp* when *δp* increases) for most parameter values. As a consequence, neglecting 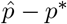 leads to:

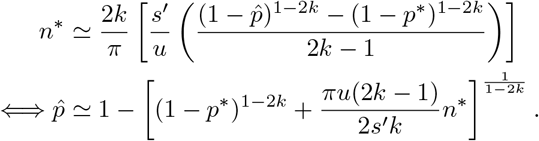

Replacing *p*\* and *n*\* with their expressions from equation 6 and reorganizing gives:

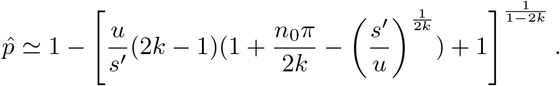

Assuming that *n*_0_ is reasonably small and *π* ≪ 1, the term *n*_0_*π*/2*k* can be further neglected, and:

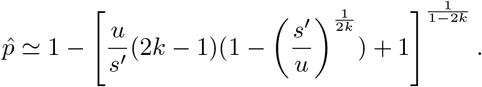

### A.5 Equilibrium stability for equation 11

The Jacobian matrix corresponding to the equilibrum 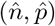 from equation 11 is:

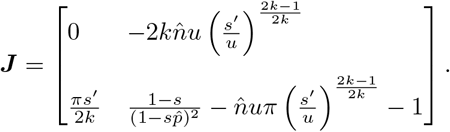

Eigenvalues are negative (i.e., the equilibrum is stable) for all tested parameter combinations. Eigenvalues happen to be complex for all parameter combinations (except for very large values of *s*), the equilibrium is thus a stable focus, reached asymptotically by oscillating around it.

**Figure A1:**
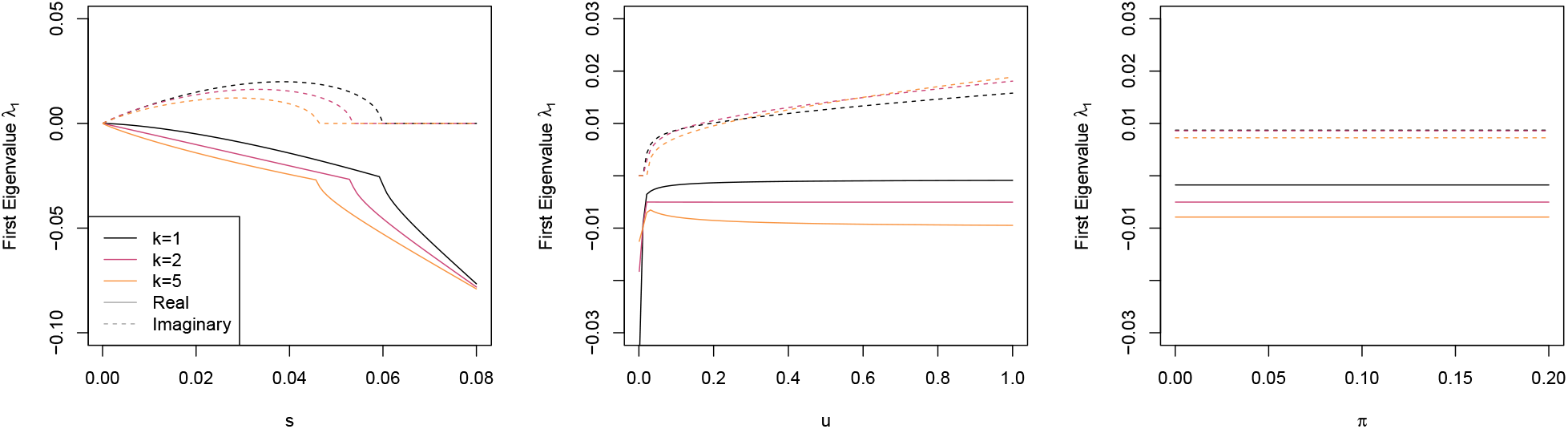
First Eigenvalue of the Jacobian matrix as a function of model parameters (*u, s*, and *π*) in the deleterious TEs - deleterious cluster model. Default parameter values were *u* = 0.1, *s* = 0.01, and *π* = 0.03. The number of clusters (*k* = 1, *k* = 2, and *k* = 5) is indicated by different line colors. Eigenvalues are complex for most of the range of the parameters, real part: plain lines, imaginary part: dotted lines.

**Figure A2:**
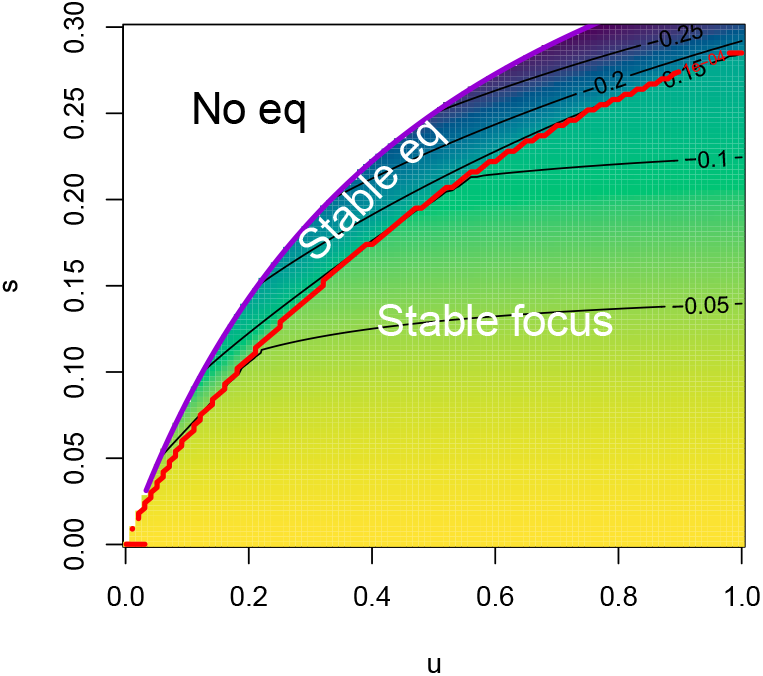
Equilibrium stability for the deleterious TEs - deleterious cluster model. The figure represents the real part of the first eigenvalue of the Jacobian matrix for two major parameters (*u* and *s*), with *k* =1 and *π* = 0.03. The eigenvalue is negative for the whole parameter range, and is a complex number for most of the range (below the red line). The purple line delineates *s* = *u*/(1 + 2*u*), beyond which selection is too strong to let the TE invade (white area).

## Appendix B Supplementary results

### B.1 Sensitivity of the neutral equilibrium (Equation 4) to model assumptions

**Figure B1:**
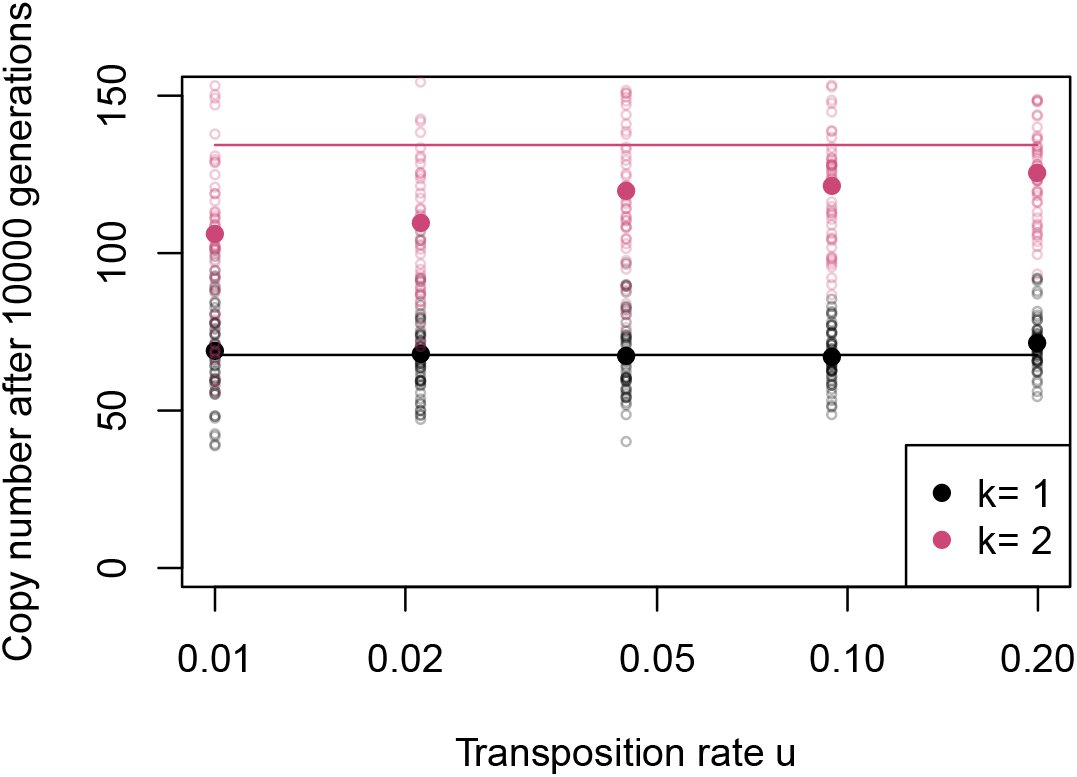
Effect of the transposition rate *u* on the simulated equilibrium copy number in the neutral model. Equation 4 predicts that, in the neutral model, the equilibrium does not depend on the transposition rate. Simulations were run for *k* = 1 and *k* = 2 clusters in populations of size *N* = 5, 000; simulations were stopped after 10, 000 generations. The figure displays the final copy number in each simulation (open symbols), their average (filled symbols), and the theoretical prediction (plain lines). Simulations display a slight increase in the equilibrium copy number for large transposition rates, due to linkage disequilibrium. This effect increases with the number of clusters. Conversely, when the transposition rate is low, the invasion dynamics is slower, and all TEs might not be fixed by the end of the simulations. Overall, theoretical predictions fit well for a single cluster, but simulations featuring several clusters are slower, and the final copy number remains below the theoretical expectation in finite populations from *k* = 2 clusters.

### B.2 Effect of genetic drift on the average and variance of the copy number

**Figure B2:**
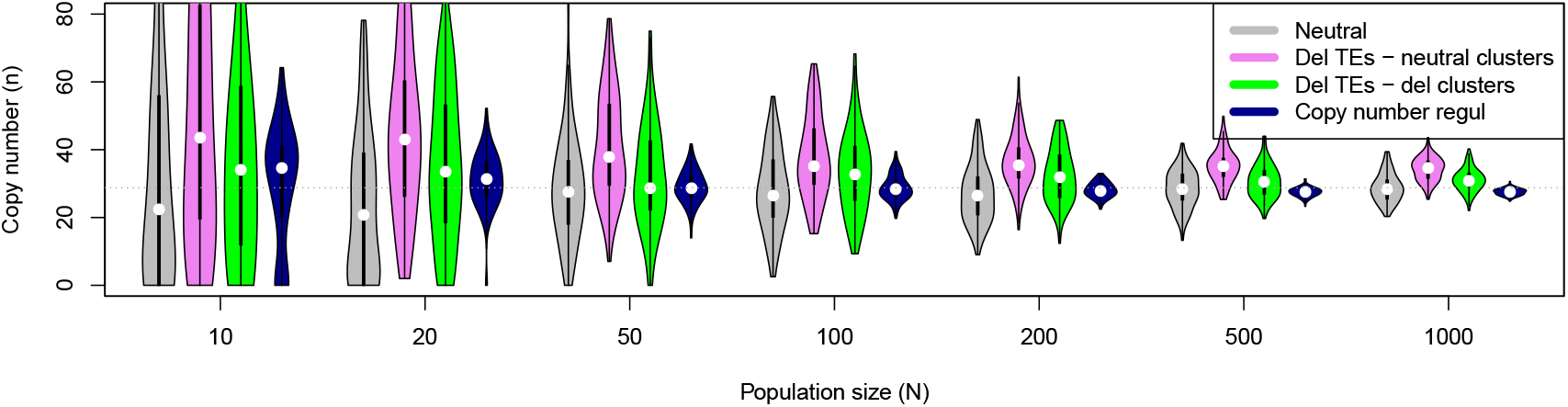
Distribution of the average copy number *n* among 1000 replicates in different models, with various population sizes. Models were parameterized so that they achieve similar average copy numbers 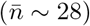 in large populations (horizontal dotted line): *s* = 0.01 for all models (except the neutral model), *k* = 1 cluster and *π* = 0.03 in all trap models. Transposition rates were: *u* = 0.045 for the neutral model, *u* = 0.05 for the Deleterious TE - neutral cluster model, *u* = 0.15 for the Deleterious TE - Deleterious cluster model, and *u_n_* = 0.17/(1 +0.45*n*) for the regulation model.

## References

Adams, M. D., R. S. Tarng, and D. C. Rio, 1997. The alternative splicing factor PSI regulates P-element third intron splicing in vivo. Genes & Development 11 (1), 129–138. doi: 10.1101/gad.11.1.129.

Arkhipova, I. and M. Meselson, 2005. Deleterious transposable elements and the extinction of asexuals. Bioessays 27 (1), 76–85. doi: 10.1002/bies.20159.

Bergman, C. M., H. Quesneville, D. Anxolabéhère, and M. Ashburner, 2006. Recurrent insertion and duplication generate networks of transposable element sequences in the *Drosophila melanogaster* genome. Genome Biology 7 (11), 1–21.

Brennecke, J., A. A. Aravin, A. Stark, et al., 2007. Discrete small RNA-generating loci as master regulators of transposon activity in Drosophila. Cell 128 (6), 1089–1103. doi: 10.1016/j.cell.2007.01.043.

Brookfield, J. F. and R. M. Badge, 1997. Population genetics models of transposable elements. Genetica 100 (1), 281–294. doi: 10.1007/978-94-011-4898-6_28.

Charlesworth, B. and D. Charlesworth, 1983. The population dynamics of transposable elements. Genetics Research 42 (1), 1–27. doi: 10.1017/S0016672300021455.

Deniz, O., J. M. Frost, and M. R. Branco, 2019. Regulation of transposable elements by DNA modifications. Nature Reviews Genetics 20 (7), 417–431. doi: 10.1038/s41576-019-0106-6.

Dolgin, E. S. and B. Charlesworth, 2006. The fate of transposable elements in asexual populations. Genetics 174 (2), 817–827. doi: 10.1534/genetics.106.060434.

Doolittle, W. F. and C. Sapienza, 1980. Selfish genes, the phenotype paradigm and genome evolution. Nature 284 (5757), 601–603. doi: 10.1038/284601a0.

Gilbert, C. and C. Feschotte, 2018. Horizontal acquisition of transposable elements and viral sequences: patterns and consequences. Current Opinion in Genetics & Development 49, 15–24. doi: 10.1016/j.gde.2018.02.007.

Gladyshev, E., 2017. Repeat-induced point mutation and other genome defense mechanisms in fungi. Microbiology Spectrum 5 (4), 5–4. doi: 10.1128/9781555819583.ch33.

Goriaux, C., E. Théron, E. Brasset, and C. Vaury, 2014. History of the discovery of a master locus producing piRNAs: the flamenco/COM locus in *Drosophila melanogaster*. Frontiers in Genetics 5, 257.

Grayling, M. J., 2014. phaseR: An R Package for Phase Plane Analysis of Autonomous ODE Systems. The R Journal 6 (2), 43–51. doi: 10.32614/RJ-2014-023.

Harris, C. R., K. J. Millman, S. J. van der Walt, et al., 2020. Array programming with NumPy. Nature 585 (7825), 357–362. doi: 10.1038/s41586-020-2649-2.

Hartl, D., E. Lozovskaya, and J. Lawrence, 1992. Nonautonomous transposable elements in prokaryotes and eukaryotes. Genetica 86 (1), 47–53. doi: 10.1007/BF00133710.

Huang, S., K. Yoshitake, and S. Asakawa, 2021. A review of discovery profiling of PlWI-Interacting RNAs and their diverse functions in Metazoans. International Journal of Molecular Sciences 22 (20), 11166. doi: 10.3390/ijms222011166.

Kelleher, E. S., R. B. Azevedo, and Y. Zheng, 2018. The evolution of small-RNA-mediated silencing of an invading transposable element. Genome Biology and Evolution 10 (11), 3038–3057. doi: 10.1093/gbe/evy218.

Khan, A. I., D. M. Dinh, D. Schneider, R. E. Lenski, and T. F. Cooper, 2011. Negative epistasis between beneficial mutations in an evolving bacterial population. Science 332 (6034), 1193–1196. doi: 10.1126/science.1203801.

Kidwell, M. G. and D. R. Lisch, 2001. Perspective: transposable elements, parasitic DNA, and genome evolution. Evolution 55 (1), 1–24. doi: 10.1111/j.0014-3820.2001.tb01268.x.

Kofler, R., 2019. Dynamics of transposable element invasions with piRNA clusters. Molecular Biology and Evolution 36 (7), 1457–1472. doi: 10.1093/molbev/msz079.

Kofler, R., 2020. piRNA clusters need a minimum size to control transposable element invasions. Genome Biology and Evolution 12 (5), 736–749. doi: 10.1093/gbe/evaa064.

Kofler, R., V. Nolte, and C. Schlötterer, 2022. The transposition rate has little influence on the plateauing level of the P-element. Molecular Biology and Evolution 39 (7), msac141. doi: 10.1093/molbev/msac141.

Kouyos, R. D., O. K. Silander, and S. Bonhoeffer, 2007. Epistasis between deleterious mutations and the evolution of recombination. Trends in Ecology & Evolution 22 (6), 308–315. doi: 10.1016/j.tree.2007.02.014.

Le Rouzic, A., 2014. Estimating directional epistasis. Frontiers in Genetics 5, 198. doi: 10.3389/fgene.2014.00198.

Le Rouzic, A. and P. Capy, 2005. The first steps of transposable elements invasion: parasitic strategy vs. genetic drift. Genetics 169 (2), 1033–1043.

Le Rouzic, A. and P. Capy, 2006. Population genetics models of competition between transposable element subfamilies. Genetics 174 (2), 785–793. doi: 10.1534/genetics.105.052241.

Lee, Y. C. G., 2022. Synergistic epistasis of the deleterious effects of transposable elements. Genetics 220 (2), iyab211. doi: 10.1093/genetics/iyab211.

Lohe, A. R. and D. L. Hartl, 1996. Autoregulation of mariner transposase activity by overproduction and dominant-negative complementation. Molecular Biology and Evolution 13 (4), 549–555. doi: 10.1093/oxfordjournals.molbev.a025615.

Lu, J. and A. G. Clark, 2010. Population dynamics of PIWI-interacting RNAs (piRNAs) and their targets in Drosophila. Genome Research 20 (2), 212–227. doi: 10.1101/gr.095406.109.

Lynch, M. and J. S. Conery, 2003. The origins of genome complexity. Science 302 (5649), 1401–1404. doi: 10.1126/science.1089370.

Maisnier-Patin, S., J. R. Roth, Å. Fredriksson, et al., 2005. Genomic buffering mitigates the effects of deleterious mutations in bacteria. Nature Genetics 37 (12), 1376–1379. doi: 10.1038/ng1676.

Malone, C. D. and G. J. Hannon, 2009. Small RNAs as guardians of the genome. Cell 136 (4), 656–668. doi: 10.1016/j.cell.2009.01.045.

Nuzhdin, S. V., 2000. “Sure facts, speculations, and open questions about the evolution of transposable element copy number”. Transposable elements and genome evolution. Springer, 129–137. doi: 10.1007/978-94-011-4156-7_15.

Orgel, L. E. and F. H. Crick, 1980. Selfish DNA: the ultimate parasite. Nature 284 (5757), 604–607. doi: 10.1038/284604a0.

Ozata, D. M., I. Gainetdinov, A. Zoch, D. O’Carroll, and P. D. Zamore, 2019. PIWI-interacting RNAs: small RNAs with big functions. Nature Reviews Genetics 20 (2), 89–108. doi: 10.1038/s41576-018-0073-3.

R Core Team, 2020. R: A language and environment for statistical computing. R Foundation for Statistical Computing. Vienna, Austria.

Robillard, E., A. Le Rouzic, Z. Zhang, P. Capy, and A. Hua-Van, 2016. Experimental evolution reveals hyperparasitic interactions among transposable elements. Proceedings of the National Academy of Sciences 113 (51), 14763–14768. doi: 10.1073/pnas.1524143113.

Roessler, K., A. Bousios, E. Meca, and B. S. Gaut, 2018. Modeling interactions between transposable elements and the plant epigenetic response: a surprising reliance on element retention. Genome Biology and Evolution 10 (3), 803–815. doi: 10.1093/gbe/evy043.

Roze, D., 2022. Causes and consequences of linkage disequilibrium among transposable elements within eukaryotic genomes. bioRxiv.

Saint-Leandre, B., P. Capy, A. Hua-Van, and J. Filée, 2020. piRNA and transposon dynamics in drosophila: A female story. Genome Biology and Evolution 12 (6), 931–947.

Selker, E. U. and J. N. Stevens, 1985. DNA methylation at asymmetric sites is associated with numerous transition mutations. Proceedings of the National Academy of Sciences 82 (23), 8114–8118. doi: 10.1073/pnas.82.23.8114.

Soetaert, K., T. Petzoldt, and R. W. Setzer, 2010. Solving differential equations in R: Package deSolve. Journal of Statistical Software 33 (9), 1–25. issn: 1548-7660. doi: 10.18637/jss.v033.i09.

Sousa, A., C. Bourgard, L. M. Wahl, and I. Gordo, 2013. Rates of transposition in *Escherichia coli*. Biology Letters 9 (6), 20130838.

Wallau, G. L., P. Capy, E. Loreto, A. Le Rouzic, and A. Hua-Van, 2016. VHICA, a new method to discriminate between vertical and horizontal transposon transfer: Application to the *mariner* family within Drosophila. Molecular Biology and Evolution 33 (4), 1094–1109.

Zanni, V., A. Eymery, M. Coiffet, et al., 2013. Distribution, evolution, and diversity of retrotransposons at the flamenco locus reflect the regulatory properties of piRNA clusters. Proceedings of the National Academy of Sciences 110 (49), 19842–19847. doi: 10.1073/pnas.1313677110.

Zhang, S., B. Pointer, and E. S. Kelleher, 2020. Rapid evolution of piRNA-mediated silencing of an invading transposable element was driven by abundant de novo mutations. Genome Research 30 (4), 566–575. doi: 10.1101/gr.251546.119.

